# Too bad to be fraud, Midjourney has yet to embark in science

**DOI:** 10.1101/2023.01.28.526052

**Authors:** Michael Cammer

## Abstract

Artificial Intelligence (AI) promises great benefits for data management, for image analysis, and for new discovery, but also creates unprecedented potential for fraud. This brief study explores whether images created by AI pose a threat to the integrity of figures in scientific publications. An attempt was made to use a leading AI image-generation engine, Midjourney, to create plausible facsimiles of images that are commonly depicted in manuscripts in the biological sciences. We conclude that a widely available AI for generated images is not a threat to integrity, yet, but may be in the near future.

## Introduction

When large annotated image datasets are used to train artificial intelligence (AI), major products may include unexpected connections, models, and image based omics. These have the potential to revolutionize science by identifying relationships and patterns that people are unable to detect which would lead to new knowledge, hypotheses, and even paradigms. The potential for such salutary outcomes is enhanced by cross referencing with other data, such as genomics, proteomics, or the wealth of literature. However, a deleterious potential is convincing simulation that could be mistaken for authentic data, even by experts (or other AIs). Science already has a Photoshop problem where people intentionally alter images to commit fraud. [Ref 1] This study explores whether there is a potential for fraud in science using a currently available popular engine for *de novo* image creation.

There are popular AI driven resources for inventing new images based on preset training. Illustrators or set designers, for instance, use them as idea factories. [ref 2] It is well established that people consider AI excellent for generating fantasy landscapes, illustrations of simple narratives, pornography, and odd juxtapositions of images that appear realistic or plausible. We chose MidJourney, a popular AI image engine running on a publicly accessible Discord server, to test whether it could generate credible scientific images.

Training sets used for AIs are critical for their output. There are libraries of microscopy images that are publicly available. For instance, the Cell Image Library [ref 3] is a repository for images with thorough tagging and cross-referenced ontologies. Although primary goals are for research, reference, and education, a long term goal is “a library of images that will serve as primary data for research.” [ref 4] A large body of images thoroughly tagged, including detailed experimental methods, could enable experimentation by query rather than by bench work; perhaps an experiment has already been done based on different inquiries or hypotheses, but a new set of eyes or analytical tools could find additional conclusions. Other groups pursue mining microscopy data for omics. [refs 5 and 6] With the advent of data repositories, such as BioImage Archive [ref 7] and Image Data Resource [ref 8], and acceleration of their use due to policies such as new NIH grant application data management plans, [ref 9] microscopy data with associated tags or linked to explanations in peer reviewed publications may become more available and useful as sources for omics and other AI mining. These data could also be used to fabricate results.

We conclude that today popular AIs are too deficient to be used for fraud in science. However, if trained by the wealth of science specific images which may already be available and which are coming online, they likely will achieve the ability to fake data.

## Methods

Text prompts were fed to the Midjourney bot via a Discord server at https://discord.gg/midjourney. Midjourney describes itself as, “an independent research lab exploring new mediums of thought and expanding the imaginative powers of the human species.” [ref 10] Most images were generated from 24 October to 6 November 2022 and additional images were generated in mid-January 2023. Each text prompt resulted in four images in a single block 512 × 512 or 1024 × 1024 pixels. All were saved as png. Simple gamma adjustment may have been applied to darker images (Photoshop 21.1.0 curves), and production of this manuscript may have reduced resolution via jpg compression unavoidable in Microsoft Word and PDF creation.

ChatGPT was used to generate answers to a query regarding microtubules (MTs) and related proteins. [ref 11]

Searches for original source materials for AI training were made in the database LAION 5B. [ref 13] This database may not be the same materials used to train Midjourney specifically, but it is indicative of how image AIs for a mass audience are being trained.

Photographs of sunflowers, zinnias, and oak were taken autumn 2022 with a Samsung Galaxy S22 cell phone. Photo of ravioli was shot circa 2007 with a Sony Cybershot camera.

Metaphase cell in Fig 5 was imaged circa 1994 with a BioRad MRC600 laser scanning confocal with Kr/Ar laser, with lines at 488, 568, and 647 nm, and a Nikon 63x N.A. 1.4 planapochromat fixed tube length lens on a Diaphot stand. HepG2 cells plated on a #1.5 coverslip were fixed with fresh PFA and labeled with fluorescence probes for beta-tubulin with a Sigma antibody (probably Cy5), f-actin with phalloidin (probably FITC), and treated with RNase and labeled for DNA with propidium iodide. Pseudocolor and other adjustments were applied in NIH-Image and Photoshop 2.5 or 3.0.

Google, Pubmed, and Wikipedia were consulted liberally.

## Summary

### Results and Discussion

MidJourney produces images based on a few words entered into the system. For instance, “/imagine landscape sunflower sunset Corot” returns, within seconds, four images. Fig 1 shows that in all four results, the AI associates Corot as oil painting and other elements of the query are illustrated explicitly and recognizably. Switching “sunflower” to “zinnia” returns images with reasonably accurate flora depicted in a style reminiscent of Jean-Baptiste-Camille Corot, the mid-Nineteenth Century French painter.

**Fig 1.**
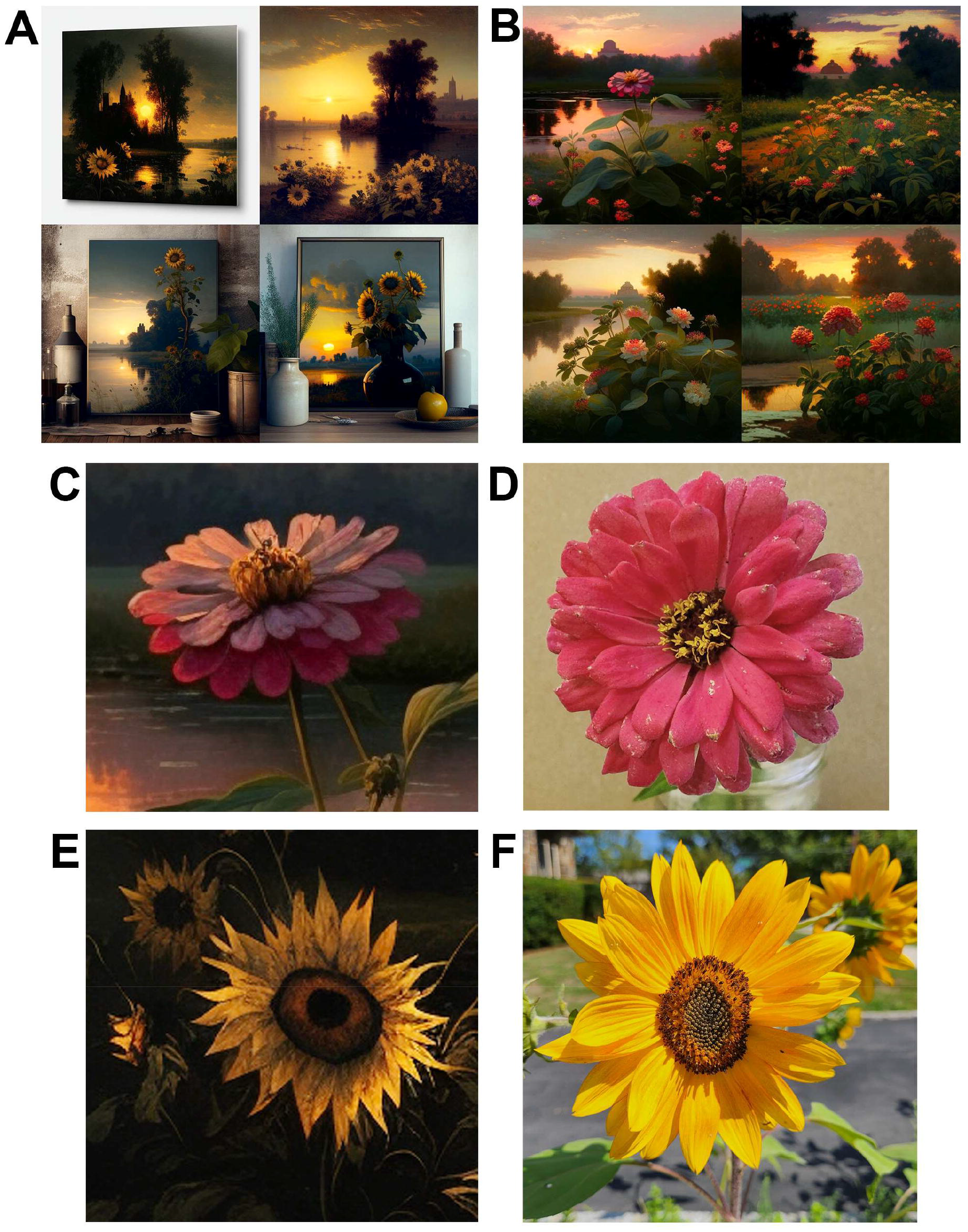
Common use of Midjourney AI to generate images in a particular style. A. Images generated with prompt “landscape sunflower sunset Corot.” B. Images generated with prompt “landscape zinnia sunset Corot.” To qualitatively assess fidelity, C-F are comparisons of AI generated zinnia (C) and photo of zinnia (D) and AI generated sunflowers (E) and sunflowers (F).

Midjourney appears to do a very good job identifying and depicting common vocabulary as judged by watching images scroll in its public interface. However, across a wide range of queries the AI returns odd results such as people with three eyes or an extra arm, misshapen animals, or other organic features which are recognizable but not constructed correctly (Supp Fig 1), but more often, it returns plausible images of animals, clothing, common objects, buildings, landscapes, and in art styles specified by the operator. How would Midjourney perform with science specific imagery?

As they are fundamental in biological science and relatively simple graphics, immunoblots are a major area of fraud in published papers [refs 18, 19, 20]. Therefore, the first scientific term tested was “western blot bands” which returned what look more like mid-20^th^ Century modern art (Figure 2) than scientific data. The lurid colors in Fig 2A resemble chromatography [ref 14] more than western blots. Constraining with terms “bands,” “lanes,” and “grayscale,” did not yield significantly more plausible results and continued to look more like modernist charcoal drawing than scientific images (Figure 2B-C). Directing the number of lanes did not work; asking for six lanes resulted in images with non-six lanes. These first tests were performed in late October 2022. Three months later, “western blot grayscale,” appeared to be retrogression (Fig 2D); the term “blot” appeared to be disassociated from a science specific context. Challenging Midjourney with “immunoblot” as a substitute for “western blot” yielded pastel paint splats and butterflies in such poor taste that they should not be displayed anywhere, even on the cheapest sweatshirt or at the kitschiest art expo marketing to hotels and strip malls. Clearly, “blot” superseded other terms and the bilateral symmetry of the ink blots probably referred to Rorschach inkblot tests specifically. Refining with “bands” as in “immunoblot scientific lanes bands” (Figure 2E) at least returned lanes of bands, flipped the contrast to represent fluorescence, but strayed into DNA motifs which seem to be the popular metonymic icon for “science.” Figure 2F is a big improvement and clearly resembles real blots such as Figure 2G, although perhaps it was trained with fluorescent immunoblots, as in Figure 2H. We conclude MidJourney is untrained in scientific western blots.

**Fig 2.**
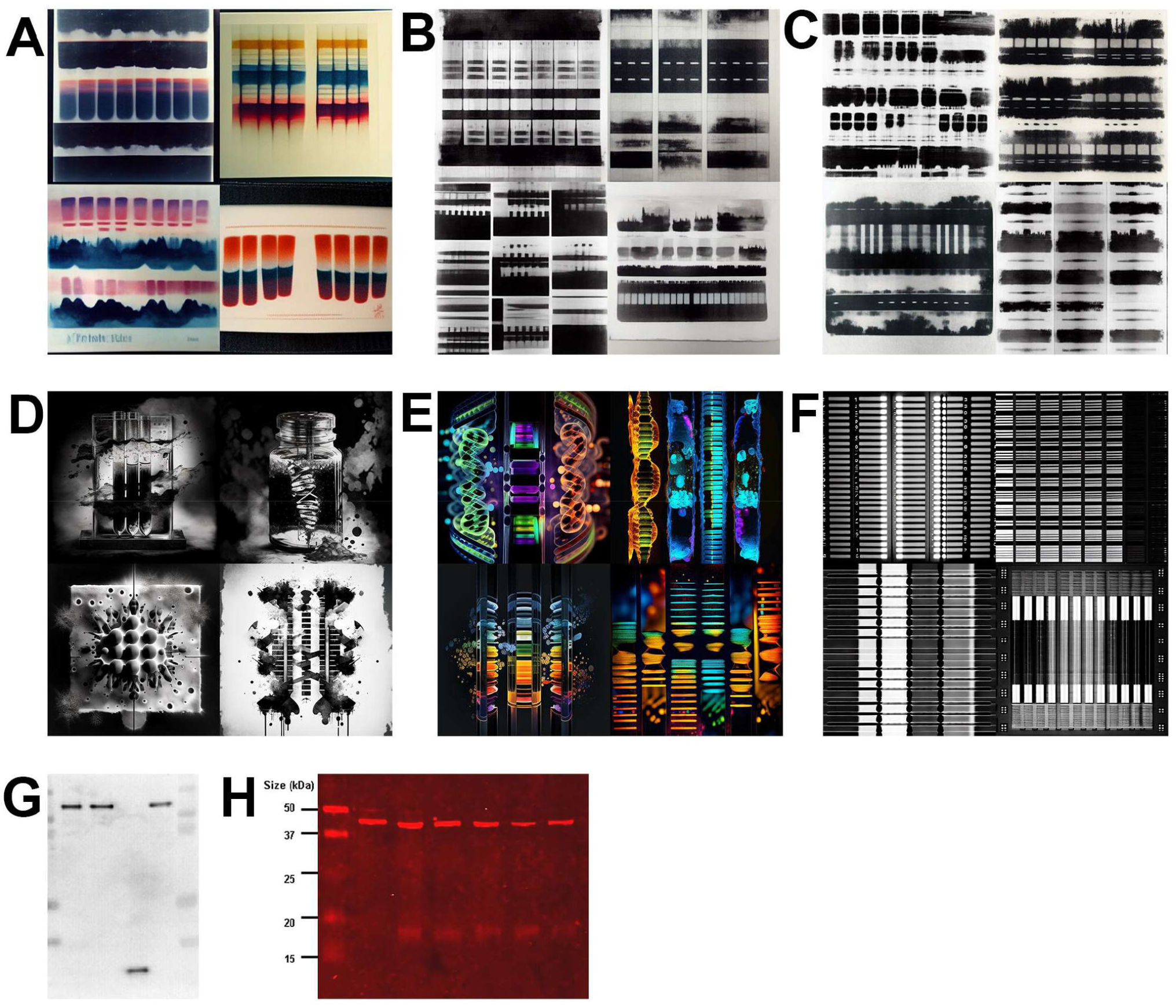
Comparison of AI generated and real western blots. A. Prompt: western blot bands. B. Prompt: western blot grayscale six lanes each lane different. C. Prompt: western blot grayscale, 6 lanes, each lane equal width, each band different, first lane 7 bands, second lane 3 bands, other lanes random bands, low noise. D: Prompt: western blot grayscale. E: Prompt: immunoblot scientific lanes bands. F: Real blot probed with anti-GFP and with molecular weight markers in partially cropped lanes at left and right margins. G: Real fluorescence immunoblot probed with anti-lipoic acid. [ref 15]

Micrographs are also fundamental in biological science and are a target for misrepresentation. We optimistically hypothesized that the AI would generate images of “cell biology microscopy.” The first result included depictions of cell monolayers, even if the details were unrealistically cartoonish (Fig 3). The AI fails at the style of micrographs, although with increased presentation of three dimensional data as highly processed reconstructions with surfaces, textures, and ray traced lighting, the style may be reasonable.

**Fig. 3.**
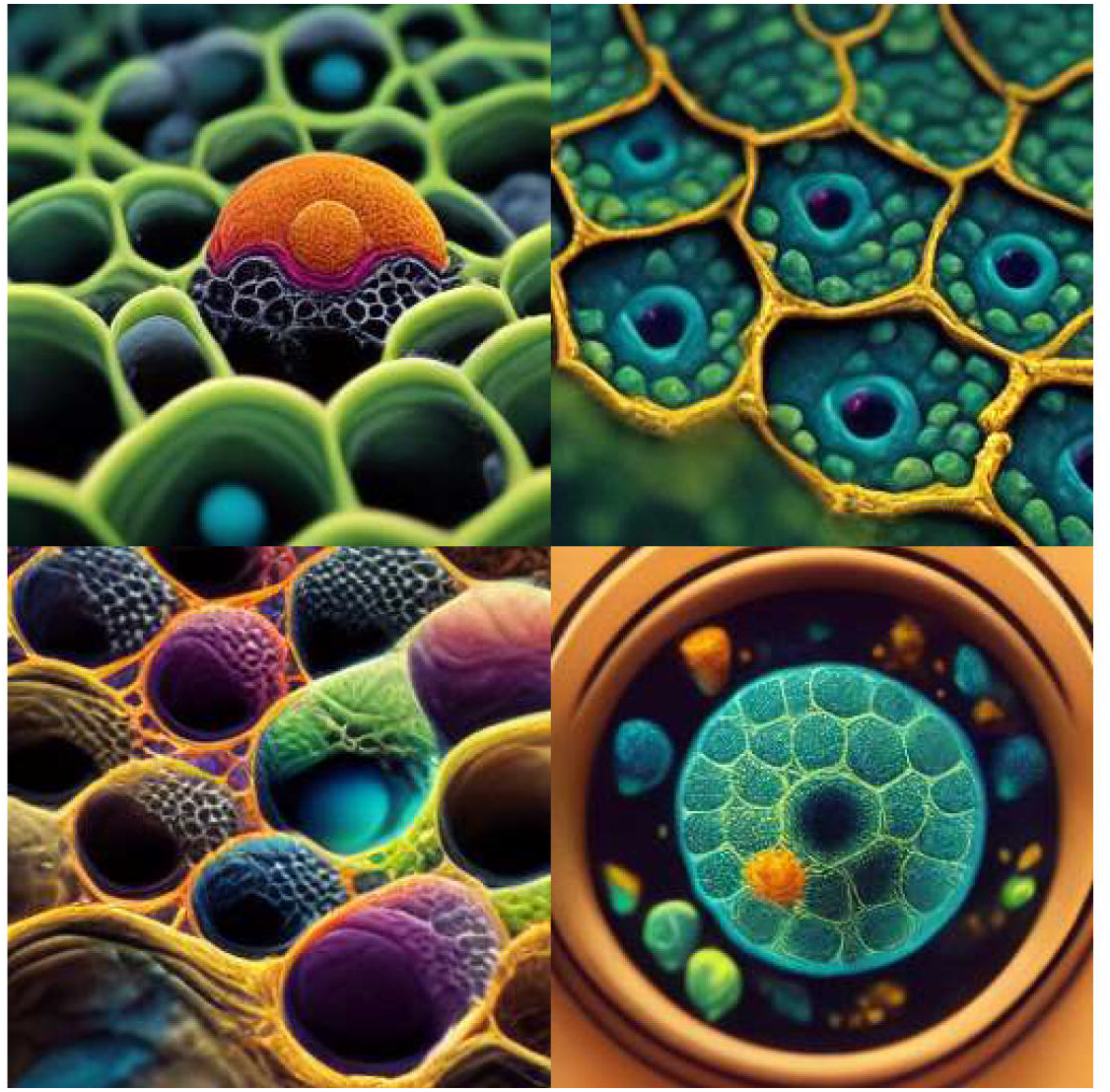
Prompt: cell biology microscopy. Whereas the two images in the left column are fanciful, the two images to the right contain basic compositions of cells in sheets, endothelia, or embryos where cell membrane were labeled. In the upper right, there is an approximation of organelles and nuclei, similar to what might be expected in a high school biology class where the assignment were to depict a monolayer of cells using different color modeling clay, although we would expect to have differentiation of Golgi and mitochondria too for an A grade.

We pushed for a more specific cell type and accurate microscopy technique presentation. In response to the prompt, **“**phase contrast macrophage microscopy fill field of view,” Midjourney produced four panels which could superficially be said to represent macrophages in a very stylized fashion (Fig. 4A), but a macrophage expert stated definitively, “That is nothing like a macrophage.” [Ref 12] Each image contains a central spot (or spots) which reflects the basic structure of a eukaryotic cell in 2D culture as having a nucleus (or nuclei). Surrounding each nucleus are elements that could be identified as highly stylized cytoplasm and organelles, but nothing identifiable as, for instance, vesicles as pictured in Fig 4B. The prompt asked specifically for phase contrast microscopy, but the lurid colors refer to either fluorescence microscopy or gratuitous use of saturated colors which appear ubiquitous throughout Midjourney’s output. Another example showing a fantastical depiction of a commonly used model organism, C. elegans, is in Supplemental Fig 2. No results resembled electron microscopy, even when explicitly prompted for this (data not shown).

**Fig. 4.**
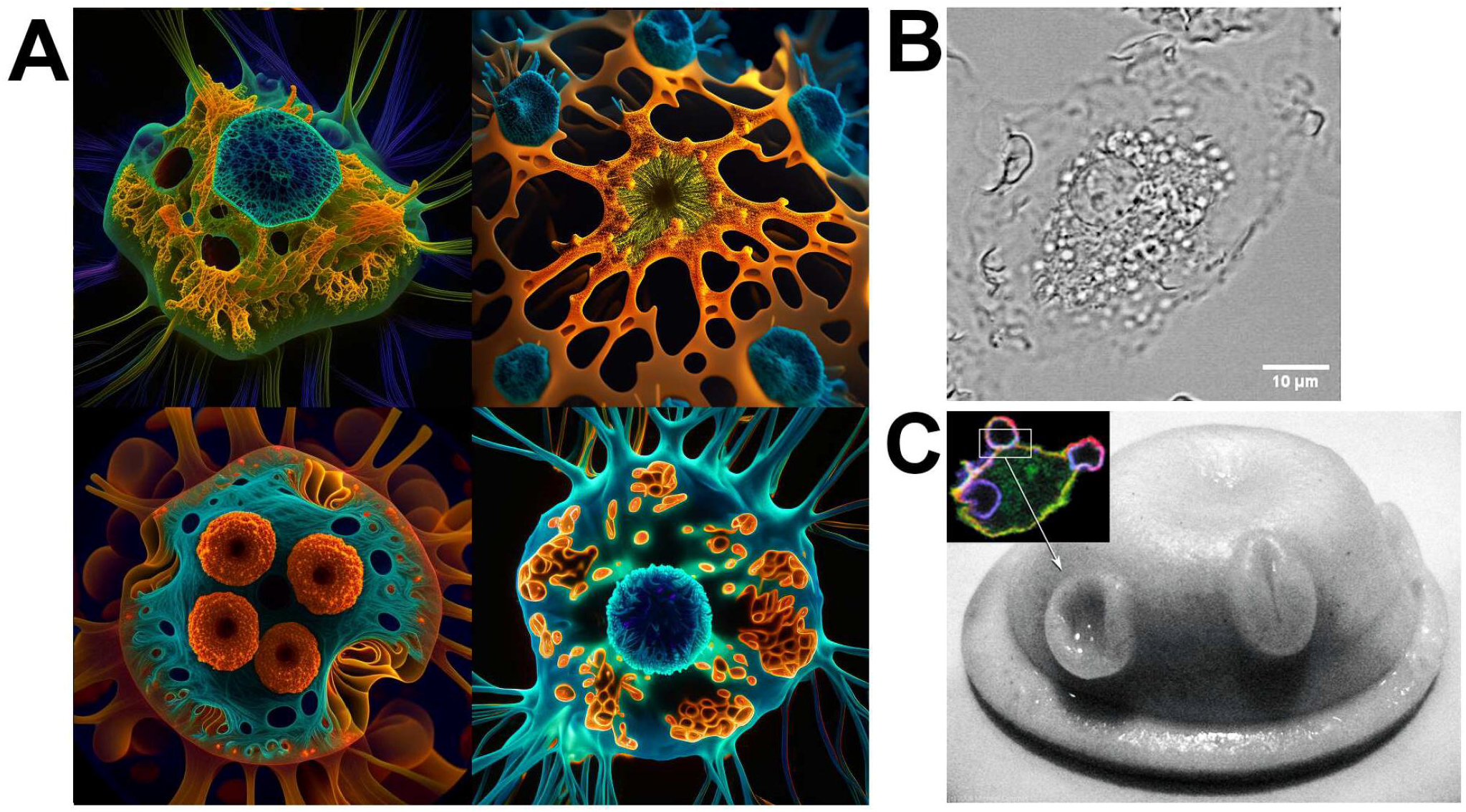
AI generated macrophage with a request of a specific imaging modality fails both to depict the cell and the contrast types. A. AI generated images with the prompt, “phase contrast macrophage microscopy fill field of view.” B. Phase contrast micrograph of live macrophage in 2D culture. (Vesicles due to LPS treatment.) C. Photo of real ravioli with popped air bubbles in the pasta shell resembles a macrophage with phagocytic cups more than AI generated images. Inset is single optical section of fluorescence stained macrophage ingesting latex beads. [ref 13] Despite bright colors on a dark background, AI images do not resemble this or other fluorescence confocal micrographs of macrophages.

Rather than pursue a cell type, we sought to reproduce a nearly ubiquitous cell biological process taught in basic biology courses as early as elementary school, eukaryotic cell division. “Micrograph of cytokinesis, tubulin is magenta, actin is cyan, cell structure multicolors,” (Fig 5A) resulted in four images where, even though pairs or grouping may refer to dividing cells, only one panel (lower left) has an obvious structure, depicted in a white outline, of a dividing cell, but the clump of four magenta circles in each daughter cell and the filamentous structures in white have no relationship to real cells. Only one panel (upper right) has any filamentous structures in cyan, and they resemble microtubules more than f-actin. In only one panel (upper right) does magenta resemble microtubules and in the lower right panel they look more like actin based structures.

**Fig. 5.**
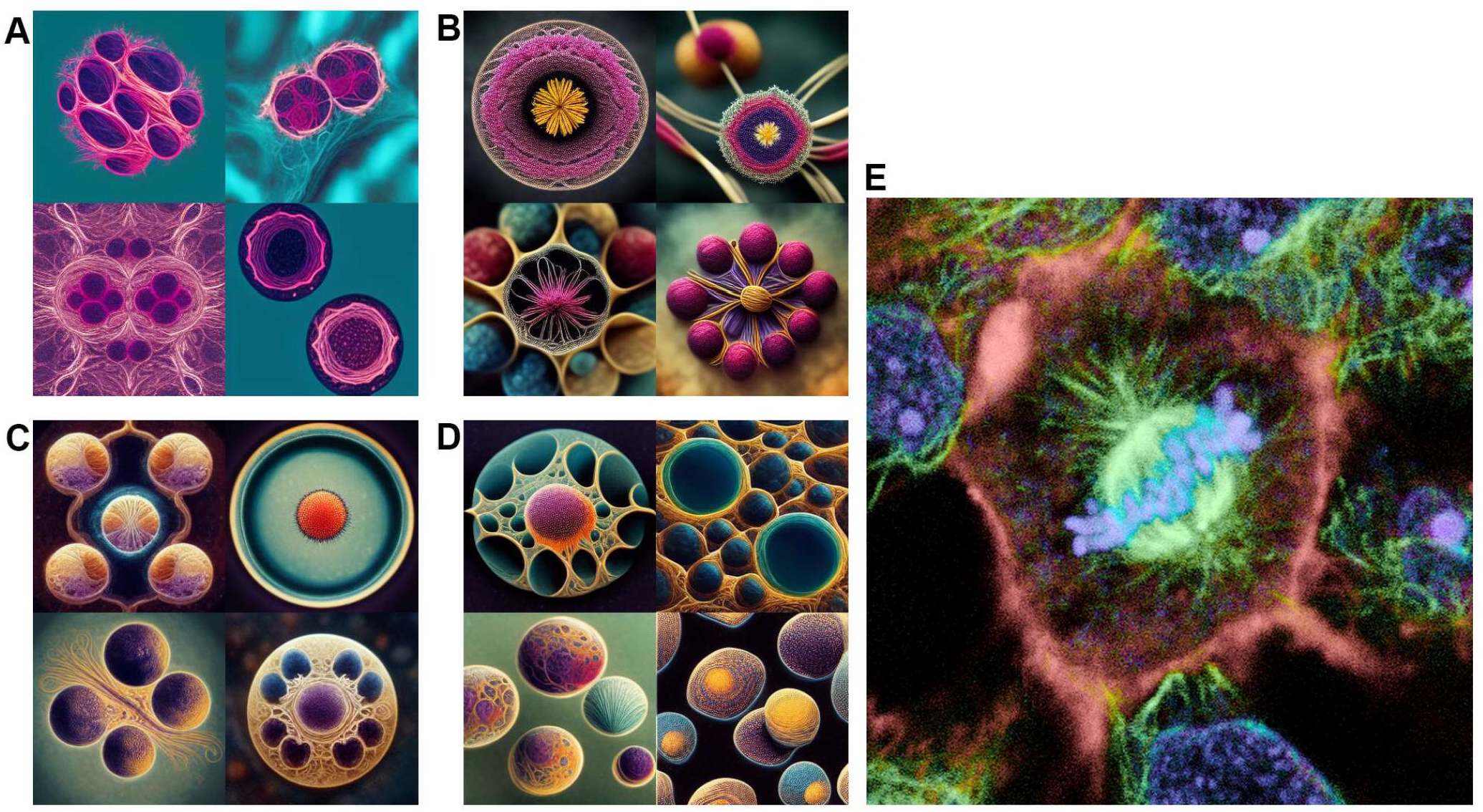
AI images of cell division do not resemble real cell division. A. Prompt: micrograph of cytokinesis, tubulin is magenta, actin is cyan, cell structure multicolors. B. Prompt: spindle, asters, chromosomes, cell division, microtubules, cell membrane, microscopy. C. Prompt: real science micrograph of cytokinesis. D. Prompt: real science micrograph of dividing cells. E. Example of real eukaryotic cell division. Laser scanning confocal optical section of fixed HepG2 cells labeled with fluorescence probes showing microtubules green, f-actin red, and DNA violet. This is an archetype of metaphase with chromosomes aligned at center of a round cell with a basket of spindles on each side and f-actin concentrated under the plasma membrane. Cell diameter approximately 30 um. None of the AI generated images resemble this or other stages of cell division. It could be argued the upper left panel of B could be asters as seen looking down at a pole of the lower right image could be interpreted as chromosomes attached to spindles, but these are generous interpretations, especially as the style of presentation in B is fiber arts.

The next failure, based on “spindle, asters, chromosomes, cell division, microtubules, cell membrane, microscopy,” is a collection of whimsical designs for fiber arts products to sell on Etsy rather than accurate representations of cytokinesis (Fig. 5B). This may be similar to artists who can make nice things with various materials but are unable to depict ideas accurately due to insufficient particular observation or knowledge.

We sought to find a middle ground between hardcore biology and common terms, a border between the popular and the scientifically particular where the AI would switch from success to failure. Basic morphology, a profile based on exterior shape, would be the primary metric for accuracy. Oak leaves have very specific morphologies that are easy to identify. The result for “oak leaf” (Fig 6A) was four large leaves depicted as autumnal brown or browning. The morphology appears to be specifically oak, but not the morphology of oaks readily available for comparison. We are too inexpert to identify the precise species of oak, but Quercus coccinea, at least to the untrained eye, looks very similar to the oaks in New York (Fig 6C) while uniquely different than the depictions in Fig 6A. The search term, “Quercus coccinea leaf,” produced essentially the same result, (Fig 6B) but not the slender less reticulated species specific ones requested. This suggests that whereas the system has been trained well enough to create a stereotypic family of a plant, it is not specifically trained in species subdivisions. (Also, for a discussion of singular “leaf” versus “leaves,” please see Supplemental Fig 1.) The upper left panel of Fig 6B also has the midrib offset where it meets the petiole which suggests the AI mimics superficial appearances without having an understanding of functional structure. This is similar to the results with cells where a general blob shape with a central circle can be generated, but no additional specifics are available. In general, the AI seems analogous to an artist untrained in science who given an assignment to depict something scientific, creates pictures based on cursory knowledge, misunderstanding, and whimsy.

**Fig 6.**
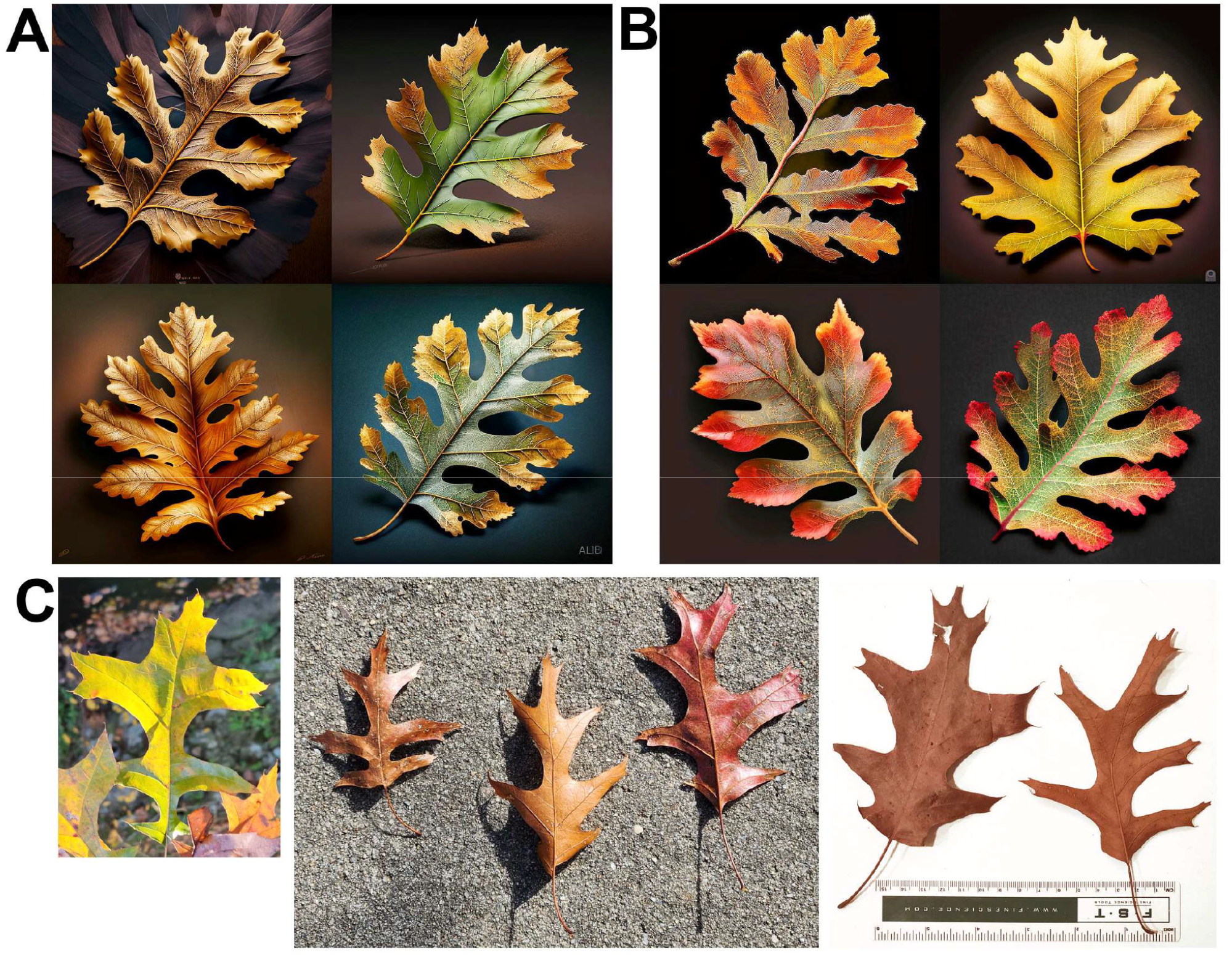
Oak leaves generated by Midjourney vs. real oak leaves. A. Prompt: oak leaf. B. Prompt: Quercus Coccinea leaf. C. Photographs of oak leaves. (A) Simple prompt “oak leaf” leads to classic story book illustrations of autumnal leaves instantly recognizable as oak. (B) Refining with specific species name does not improve quality. The AI recognizes the reference to oak, but not the specific morphology of the variety. (C) Photographs of oak leaves in New Rochelle and Larchmont, NY in October 2022 have fewer crenulations which is specific to their species.

Perhaps images generated by the database do not resemble real micrographs because it has been populated with drawings and models, not direct images of real phenomena. Reviewing hits for “microtubules” in the database LAION 5B, which was used for training Stable Diffusion, a similar engine to Midjourney, we found that most images were models, not micrographs.

The situation appears to be far more mature with text based AI. As the following interaction with ChatGPT shows, whereas Midjourney’s constructions when confronted with scientific terms are risible, ChatGPT may be ready to teach cell biology (or replace microscopy core staff). [ref 11]

> Query: Please tell me about proteins that translocate on MTs
>
> ChatGPT: Proteins that translocate on microtubules (MTs) are a diverse group of molecules that have the ability to move along the length of these cytoskeletal structures. Some examples of proteins that translocate on MTs include kinesin, dynein, and myosin. These proteins bind to MTs and use the energy from ATP hydrolysis to move along the MTs in a specific direction. Kinesin is an example of a protein that moves towards the plus end of the MT, while dynein moves towards the minus end. Myosin is another protein that can move along microtubules. These proteins play important roles in various cellular processes such as cell division, intracellular transport, and cell motility.
>
> Query: Do you know which statement of fact in your answer is incorrect?
>
> ChatGPT: My apologies, I made an error in my previous response. Myosin is a motor protein that binds to actin filaments and uses the energy from ATP hydrolysis to generate force for muscle contraction and other cellular movements, it does not move along Microtubules.

ChatGPT apparently has much deeper training. ChatGPT has resulted in a plethora of discussions regarding how effectively it mimics human writing and what implications there are (because this is so ubiquitous in popular and social media, I am not providing specific references). The wrong answer, while wrong, sounded reasonable. When identified, there was sufficient material in the data to correct the error. (And unlike real scientists, the correction was direct and unimpeded by ego issues.) This is far more sophisticated than Midjourney’s fabulous visual constructions. We attribute this to insufficient data. Imagine the possibilities when there is sufficient data, such as when repositories required by funding agencies are populated with well tagged images. There are also treasure troves, silos, and locked boxes, servers, and stacks of hard drives, packed with data which, if tagged and available could provide a wealth of material for AIs.

At this point, publicly available AI image tools are not prepared to fake science images because both the verbal and visual vocabularies of science are missing. On the other hand, training with large numbers of properly tagged images, which would be valuable for discoveries of patterns not yet realized by humans, would lead to the ability to synthesize fake data.

## Acknowledgements

Original microscopy performed at Analytical Imaging Facility of Albert Einstein of Medicine and at NYU Langone Medical Center Microscopy Core. The former is partially funded by cancer center support grant NCI P30CA013330 and the latter by NCI P30CA016087. Metaphase sample prepared by Dr. Phyllis Novikoff. Special thanks to Dr. Dianne Cox for western blot in Figure 2 and support throughout this project. Original concept for this manuscript can be found at http://microscopynotes.com/AI/01denovo/index.html.

---

**Supp Fig 1.** AI has difficulty with plurals and structure versus superficial appearance.

A. Prompt: quercus coccinea leaves.

B. Prompt: ground covered with anatomically correct fallen oak leaves

(A) Plural “leaves” depicts some stem and leaf morphology accurately but also jumbles morphology. Except in the lower left panel, species specific information lost. Similarly, does the zinnia in Fig 1C have two stems? It is also interesting that the prompt was unspecific about the context of the leaves. All results were depicted on face, late season, and against a dark background. A carpet of fallen leaves on the ground or a tree full of leaves would have been alternative acceptable results. Thus the prompt (B) was given which resulted in the upper left panel’s double apex leaf and the upper right panel’s branched triple apex leaf. Much biology is very specific about numbers and functional structural connections which are violated in these results. Also, contrary to the demand for oak specifically, non-oak leaves were included.

**Supp Fig 2.** AI generated C. elegans images do not resemble real worms.

A. Prompt: c elegans.

B. Prompt: c elegans microscopy.

C. Typical live adult C elegans imaged by Nomarski contrast. [Ref 17] (A-B) Midjourney generated images of C elegans stray from the simple stereotypical structure of real worms. High intensity and saturated colors on a black background may draw from fluorescence microscopy source material, but do not reproduce typical colors in scientific literature and fail to converge on basic worm morphology. Not having spatial scale bars is the least concern.

