## Supplementary figures and images for "Too bad to be fraud, Midjourney has yet to embark in science"

### Supplemental Fig 1

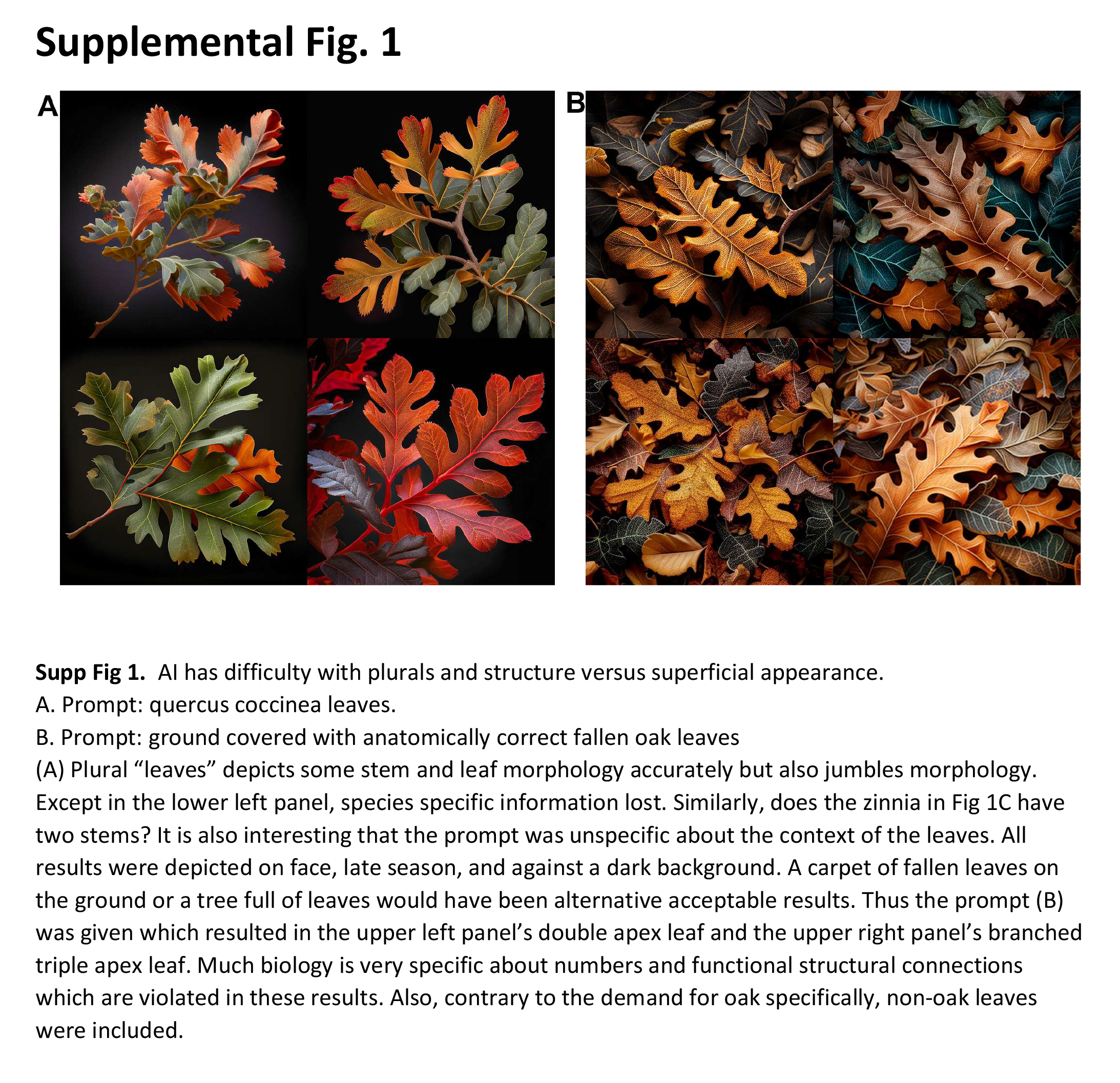

### Supplemental Fig 2

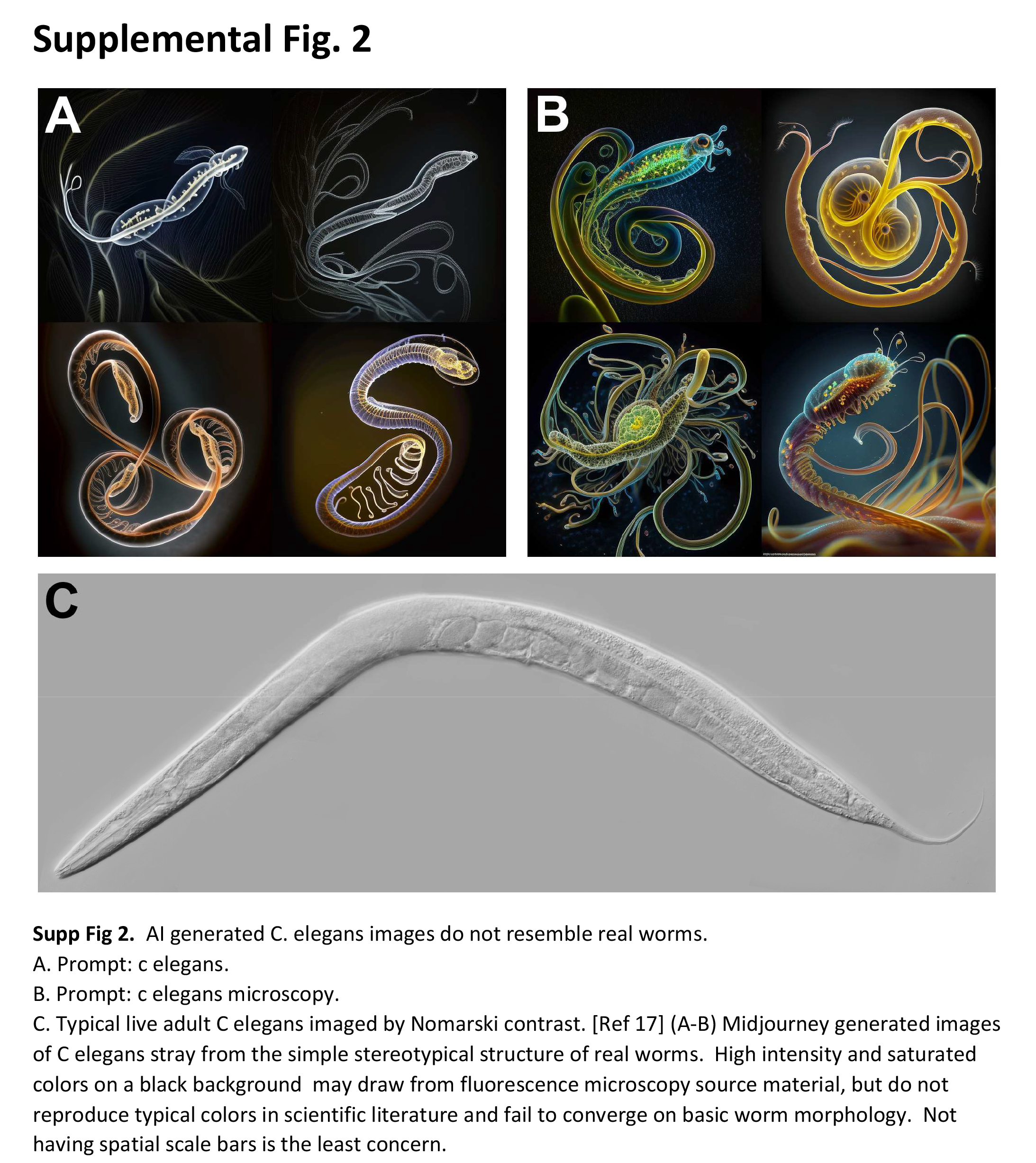
